# Calcitriol impairs the secretion of IL-4 and IL-13 in Th2 cells via modulating VDR-Gata3-Gfi1 axis

**DOI:** 10.1101/2023.12.03.569823

**Authors:** Biswajit Biswas, Shagnik Chattopadhyay, Sayantee Hazra, Ritobrata Goswami

## Abstract

Calcitriol, the bioactive form of vitamin D, exerts its biological functions by binding to its cognate receptor VDR. The indicators of the severity of allergies and asthma have been linked to low vitamin D levels. However, the role of calcitriol in regulating IL-4 and IL-13; two cytokines pivotal to allergic inflammation, remained unclear. Our study observed a decrease in IL-4 and IL-13 secretion in murine and human Th2 cells, treated with calcitriol. In murine Th2 cells, Gata3 expression was attenuated by calcitriol. However, the expression of the transcriptional repressor Gfi1, that prevents Th1 fate of activated CD4+ T cells, too was attenuated in the presence of calcitriol. Ectopic expression of either Gfi1 or VDR impaired the secretion of IL-13 in Th2 cells. Gfi1 significantly impaired *Il13* promoter activation; which calcitriol failed to restore. Ecr, a conserved region between the two genes, which enhanced the transactivation of *Il4* and *Il13* promoters; is essential for calcitriol-mediated suppression of both the genes. In murine Th2 cells, VDR interacted with Gata3 but not Gfi1. Calcitriol augmented the recruitment of VDR to the *Il13* promoter and Ecr regions. Gata3 recruitment was significantly impaired at *Il13* and Ecr locus in the presence of calcitriol but increased at *Il4* promoter. However, in the presence of calcitriol, the overall recruitment of Gfi1 remained unchanged at *Il4* but increased at *Il13* locus in Th2 cells. Together, our study clearly elucidates that calcitriol modulates VDR, Gata3, and Gfi1 to suppress IL-4 and IL-13 production in Th2 cells.

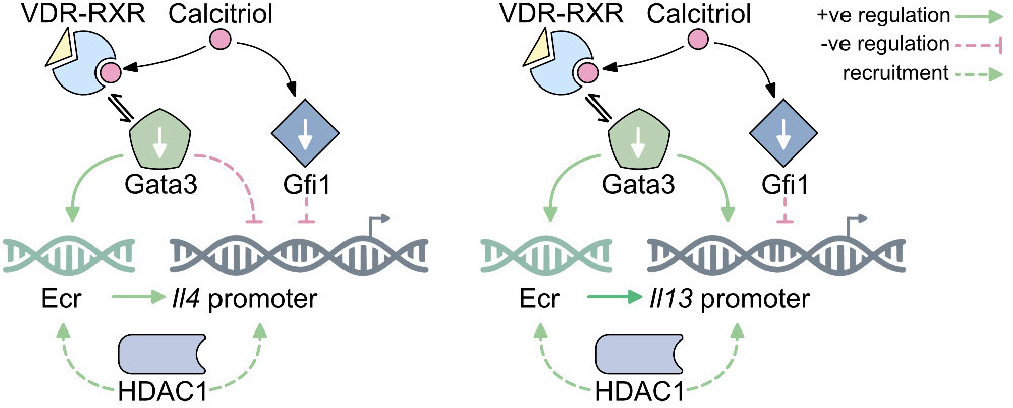

## Introduction

Originally it was observed that upon encountering pathogens, naïve CD4+ T cells would differentiate primarily into two subtypes, Th1 and Th2, determined by the cytokines present in the microenvironment (1). However, the entire gamut of T helper cell phenotype altered following the discovery of Th17, and Treg cells. Further studies unraveled the discovery of Th9 cells, Tfh cells, and Th22 cells, that fine-tunes the regulation of adaptive immune system from various pathogenic insults (2). The key cytokines produced by Th2 cells are IL-4, IL-5, IL-13, and IL-10. Th2 cells would aid in B cell activation and antibody production, especially IgE secretion (1). IL-4 and IL-13 are known to drive IgE production and effector functions of mast cells, which is directly associated with atopic diseases (3). Dysregulated Th2 cells are the main drivers of allergic asthma and rhinitis (4). The inflammatory responses, led by excess mucus production and smooth muscle contraction in the respiratory tract, are predominantly mediated by IL-13, with IL-4 playing second fiddle (5, 6). The co-expression of IL-4 and IL-13 in Th2 cells results in the convergence of their roles, leading to significant functional overlap (7). *Il4* and *Il13* genes are found in close proximity in the vertebrate genome, and share many cis-regulatory elements (8), indicating they might have emerged from a gene duplication event at some point in vertebrate evolution (7). A distantly placed regulatory region between the *Il4* and *Il13* genes with a strong affinity for Gata3 was mentioned in one of the earlier studies (9). This element was proved to be an enhancer for *Il4* (9). However, its role in *Il13* regulation has never been evaluated.

Recent studies suggest that calcitriol might have a crucial role in the preservation of immunological balance by controlling various immune cells (10–12). Upon binding to VDR, calcitriol forms a heterodimer with RXR (retinoid X receptor), and translocate to the nucleus (13). Being promiscuous in nature, calcitriol-bound VDR can form complexes with other transcription factors, hence significantly augmenting its capacity to regulate a wide array of genes (13, 14). In one of our previous studies, we observed calcitriol to be a negative regulator of Th9 phenotype (15).

Studies have linked asthma and allergic rhinitis with calcitriol deficiency (16, 17). However, no concrete association could properly be established. In psoriasis patients, calcitriol ointment provides a great deal of versatility in terms of usage in different monotherapy and combination therapy regimens (18). In contrast to common observations regarding calcitriol being a positive regulator for type 2 responses, a study in 1994 by Freedman and his group had shown that calcitriol attenuated *Il4* expression during in vitro polarization of murine CD4+ T cells (11, 19). However, the majority of the studies lack molecular mechanisms behind calcitriol-mediated regulation of Th2 phenotype. VDR could naturally inhibit IL-13 in esophageal epithelial cells (20). Therefore, it is imperative to elucidate the role of calcitriol in the regulation of Th2 cells.

Our study demonstrated an impaired production of IL-4 and IL-13 in Th2 cells treated with calcitriol. Gata3 and Gfi1, two essential transcriptional regulators of Th2 cell development, were down-regulated by calcitriol. When ectopically expressed, Gfi1 inhibited Th2 cells’ ability to produce IL-13. Both *Il4* and *Il13* genes were induced by the same regulatory element, Ecr, without which the repression of *Il4* and *Il13* genes would not have happened. At the *Il4*/*Il13* locus, calcitriol changed the recruitment of transcription factors (Gata3, Gfi1), histone modifier (HDAC1), and a specific histone modification (AcH4K5). Overall, our research underscores the function of calcitriol in Th2 cell development, with an emphasis on the underlying molecular pathways.

## Materials and methods

### Source of T cells

Naïve CD4+ T cells were isolated from wild-type C57BL/6 mice aged between 8–10 weeks. The mice used in this study were obtained from the National Institute of Nutrition in Hyderabad, India. The subjects were accommodated and propagated within the animal housing facility, Indian Institute of Technology Kharagpur, under sterile and pathogen-free conditions.

Human PBMC (peripheral blood mononuclear cells) were isolated from healthy volunteers. First, blood was drawn with the help of a phlebotomist and diluted with PBS in a 1:1 ratio. Then the PBMC were separated by density gradient centrifugation using Histopaque (Sigma-Aldrich, St. Louis, MO, USA) and collected in a different tube. All the studies conducted in this research conformed to the ethical guidelines of the Institute Animal Ethics Committee of Indian Institute of Technology Kharagpur.

### Primary cell culture

1x10^6^ cells/mL of Naïve CD4+ T cells were isolated from murine spleen and lymph-nodes via magnetic cell sorting (MojoSort, BioLegend, San Diego, CA, USA; #480039). Human Naïve CD4+ T cells were isolated (1x10^6^ cells/mL) from PBMC, using MojoSort isolation kit for humans (#480041). After isolation, naive CD4 + T cells were suspended in complete RPMI-1640 (supplemented with 10% FBS and 1% Penicillin-Streptomycin) and activated in the presence of antibodies against CD3 (145-2C11, 2 µg/mL) and CD28 (37.51, 1 µg/mL) (15). Naïve CD4+ T cells were polarized under Th2 differentiating conditions using recombinant IL-4 (20 ng/mL) and neutralizing IFN-*γ* (XMG1.2, 10 µg/mL) for 5 days with an expansion on day three (15). Calcitriol used in all experiments was procured from Selleck Chemicals, Houston, TX, USA. Unless mentioned otherwise, 100 nM calcitriol was used during differentiation. The viability of the Th2 cells upon calcitriol treatment was determined by MTT assay (10, 25, 50, and 100 nM calcitriol) and DAPI (100 nM calcitriol) staining (data not shown). In certain experiments, Th2 cells were treated with trichostatin A (TSA, 10 ng/mL) (Sigma-Aldrich, St. Louis, MO, USA) for 5 days of differentiation.

### Cloning

The pGL3-basic and pRL-TK vectors were a kind gift from Mark H Kaplan, Indiana University, USA. The murine *Il4* and *Il13* promoters were cloned from the murine genomic DNA. The constructs were sequence verified and subsequently used in all downstream luciferase assays. Likewise, the 649 base-pair cECR fragment was also isolated from the genomic DNA, inserted upstream of the *Il4* and *Il13* promoters **(Figure 6)** and sequence verified.

### Flow cytometry and intercellular cytokine staining

After 5 days of culture, differentiated Th2 cells were restimulated with anti-mouse CD3 (4 µg/mL) for 3 hours followed by 2 hours of incubation with monensin (Thermo Fisher Scientific, MA, USA) at 37 °C. Cells were surface stained with anti-mouse CD4 antibody (BioLegend) and fixed with 2% formaldehyde (HiMedia Laboratories, Maharashtra, India). Cells were then permeabilized with permeabilization buffer and stained using fluorochrome conjugated anti-mouse IL-4-APC (11B11, #504106), IL-13-PE/Cy7 (eBio13A, #25-7133-80), IL-10-FITC/PE (JES5-16E3, #11-7101-82, and #505007), IL-9-PE (RM9A4, #514104), andIFN-*γ*-PE (XMG1.2, #505808) antibodies. As a selection marker, CD4-FITC (#100406) was used. Anti-mouse IL-10 and anti-mouse IL-13 antibodies were purchased from Invitrogen, Thermo Fisher Scientific; and the remaining antibodies were procured from BioLegend. Flow cytometry experiments were performed in LSR Fortessa and FACSCalibur (BD Biosciences, San Jose, CA, USA). The data were analyzed with FCS Express 7, De Novo Software, CA, USA.

### Real-time PCR

On day 5, Th2 cells in the presence or absence of calcitriol were re-stimulated with 4 µg/mL anti-CD3 antibody for 6 hours. Post stimulation, total RNA was isolated with TRIzol (Thermo Fisher Scientific). cDNA was synthesized with iScript cDNA synthesis kit (Bio-Rad, Hercules, CA, USA), and real-time PCR was performed with PowerUp SYBR Green PCR Master Mix in QuantStudio 5 system (Thermo Fisher Scientific). Relative fold change was calculated with 2^−ΔΔ*Ct*^ methods, and plotted as individual values with a bar representing mean and an error bar representing SEM (standard error of mean) in GraphPad Prism 9 for windows (GraphPad Software, CA, USA). The sequences of the primers have been provided in Table 1. Additional primer information is already available from our previous studies (15, 21).

**Table 1.**
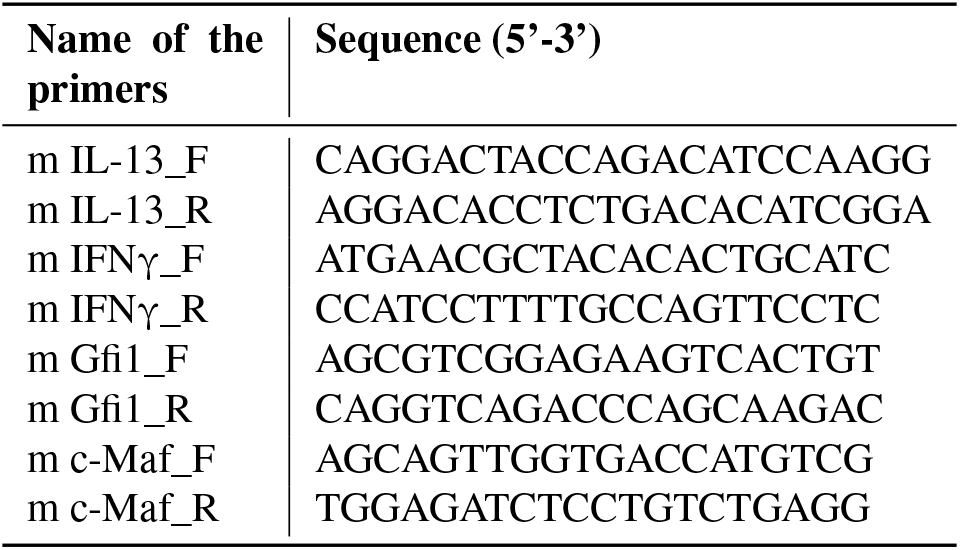
qRT-PCR primer list.

### Enzyme-Linked Immunosorbent Assay

5-day differentiated cells were re-stimulated with anti-mouse CD3 (2 µg/mL) for an additional 24 hrs. Post stimulation, the media was collected and used as samples. Capture and biotin conjugated detection anti-mouse IL-4 antibodies were purchased from BioLegend. Anti-mouse IL-13 antibody pair were purchased from Invitrogen. Standards for both IL-4 and IL-13 were purchased from BioLegend. IL-10 ELISA kit was purchased from BioLegend.

### Overexpression studies

MIEG-mVDR-hCD4, and MSCV-mGfi1-IRES-GFP (#91891, Addgene, MA, USA) were co-transfected with viral envelope constructs Gag-Pol and Env into HEK293T cells by calcium-phosphate transfection method. The supernatant with retroviral particles was harvested after 48 hours. On day 2 of Th2 cell culture, the differentiating cells were transduced with mVDR and Gfi1-expressing retrovirus particles in the presence of 10 µg/mL of Polybrene transfection reagent (Merck Millipore, Burlington, MA, USA). These cells were then subsequently expanded on day 3 and finally harvested for flow cytometry on day 5. In the mVDR over-expression studies, calcitriol has been added to the culture medium in day 3 as the ligand of VDR. In all the over-expression studies, anti-mouse CD4-PE/FITC (#100511 and #100406) were used as selection markers. In mVDR study, anti-human CD4-PE-Cy7/APC (#557852 and #17-0048-41) were used to asses transduction efficiency. The gating strategy of mGFI1-GFP is shown in **Figure 4A**. The isotype controls are shown in **Supplementary figure 1B,C**. In all the experiments, the empty vectors without mVDR (MIEG-hCD4) or Gfi1 (MSCV-IRES-GFP) constructs worked as controls.

### Immunoblotting and Immunoprecipitation

In the presence or absence of calcitriol, total cell lysates were prepared using RIPA buffer from differentiated Th2 cells. The lysates were resuspended in Laemmli buffer and boiled at 100°C for 10 mins, run on SDS-PAGE gel, transferred on PVDF membrane and probed with primary antibodies [VDR (#12550S), Gata3 (#5852S), Gfi1 (#31929S), and PU.1 (#2266S); Cell Signaling Technology (CST), Danvers, MA, USA]. The β-Actin (#AC026) and the HRP-conjugated anti-rabbit IgG (#AS014) secondary antibodies were purchased from Abclonal Technology, Woburn, MA, USA. The membrane was developed with ECL reagent (Takara). Whole cell lysates of Th2 cells were prepared and immunoprecipitated with 2 µg of pulldown antibodies (Gata3, VDR, Gfi1) using protein G agarose beads (Santa Cruz Biotechnology, Dallas, TX, USA). Eluted conjugates were run on SDS-PAGE and probed with primary antibodies of partner proteins. The membrane was then incubated with Protein-A-HRP conjugate (#12291S, CST) and developed with ECL reagents.

### Reporter assay

Dual-luciferase reporter assay was performed to assess regulatory functions of *Il4* and *Il13* promoters along with the Ecr region (previously described in cloning section). HEK293T cells (kind gift from Dr. Arnab Gupta, IISER Kolkata) were transfected (Xfect transfection reagent, Takara, Kusatsu, Japan) with p4-pGL3, p13-pGL3, p4-Ecr-pGL3, and p13-Ecr-pGL3 vectors respectively, in different combinations. To assess the role of Gata3 and Gfi1, cells were co-transfected with pcDNA-mGATA3 (#1332, Addgene) and MSCV-mGfi1-IRES-GFP expression plasmids, in the presence or absence of calcitriol. The mVDR mutations (ΔDBD, Δ403, and Δ 276) were previously generated in our lab, and were used to assess possible roles of different mVDR domains in regulating *Il4* and *Il13* promoters. pRL-TK was used as internal control in 1: 100 ratios with pGL3-based vectors.

### Chromatin immunoprecipitation

Chromatin immunoprecipitation assay was performed with Th2 cell as described previously (15). Briefly, Th2 cells treated with or without calcitriol were fixed and DNA-protein were crosslinked after 5 days of differentiation. Subsequently, the cells were sonicated to generate fragments of chromatin. The following antibodies [VDR, Gfi1, Gata3, AcH3K9 (#9649T), AcH4K5 (#9672S), and HDAC1 (#34589S)] were purchased from CST and used for immunoprecipitation. Following the reversal of the cross-links, the corresponding DNA fragments are eluted, and analyzed by quantitative RT-PCR (qRT-PCR). To determine the percentage of input, qRT-PCR was carried out using the precipitated chromatin. The data were then normalized using the appropriate isotype control, and the percentage of input chromatin was determined. All the used primers have been listed in Table 2.

**Table 2.**
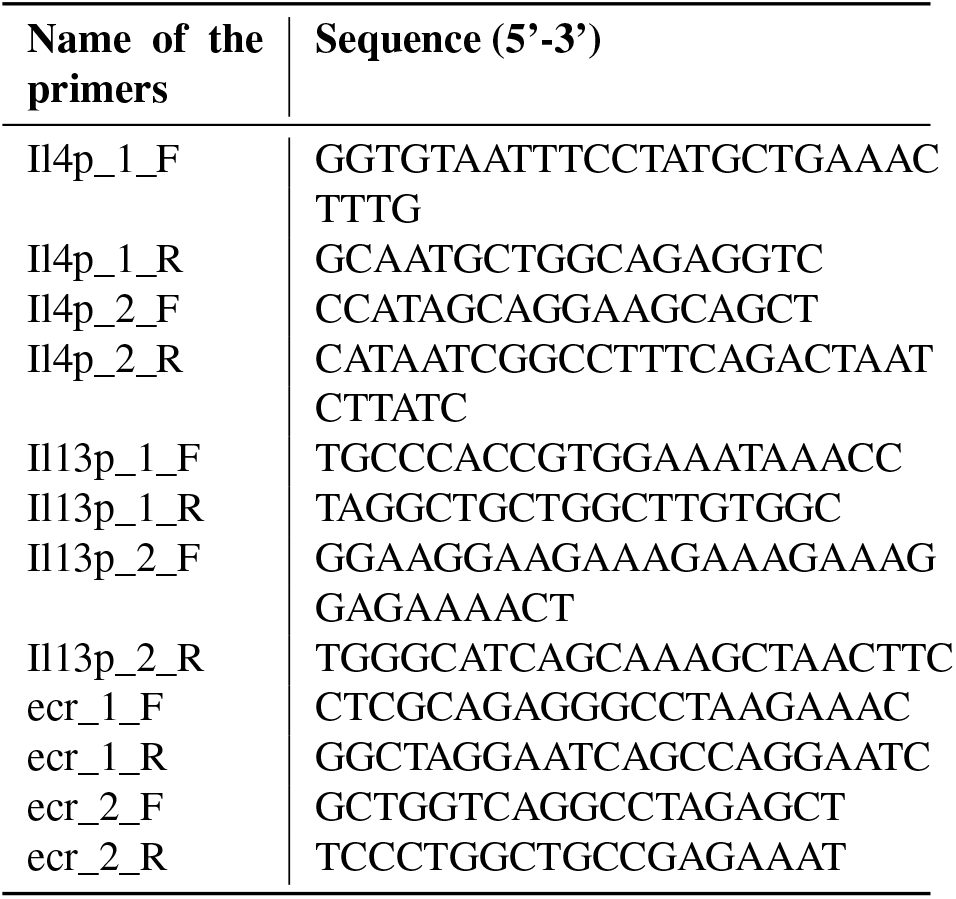
ChIP primer list.

### Statistical analysis

To identify the difference between two groups, unpaired parametric two-tailed t-test was used, assuming both populations have equal standard deviations (SDs). Ordinary one-way ANOVA followed by Tukey’s multiple comparison test was performed assuming equal SDs, to calculate the levels of variation between two or more groups. To assesses the difference in grouped data, ordinary two-way ANOVA was performed, followed by Tukey’s multiple comparison test, with a single pooled variance. In graphs, the bars represent mean with the error bars represent standard error of mean (SEM). Confidence was set to 95% with p value *<*0.05. *p*<*0.05, **p*<*0.01, ***p*<*0.001, ****p*<*0.0001, ns = not significant.

## Results

### Calcitriol suppressed IL-4 and IL-13, but induced IL-10 production in Th2 cells

VDR is the high affinity receptor for calcitriol, and is a nuclear receptor by nature.We observed an increase in VDR expression in calcitriol-treated Th2 cells **(Figure 1A)**. IL-4 and IL-13 are the two major type 2 cytokines produced by Th2 cells. *Il4* and *Il13* genes are proximally located in the vertebrate genome, have overlapping functions and share common receptors. Hence, any dysregulation in IL-4 and IL-13 production can lead to the onset of allergic diseases (3). Th2 cells also produce IL-10 in a large quantity, only second to Treg cells (22). IL-10 usually functions as an anti-inflammatory cytokine (23). In our study, we found that calcitriol reduced the percentage of IL-4- and IL-13-positive Th2 cells in a dose-dependent manner **(Figure 1B, C)**. We observed 100 nM being the most effective dose **(Figure 1D)**. Hence, we used 100 nM concentration of calcitriol in all subsequent experiments. 100 nM calcitriol is equivalent to 41.6 ng/mL, which is well within the recommended level in a healthy individual (24). We also found decreased expression of *Il4* and *Il13* in murine Th2 cells treated with calcitriol, which is in concert with the observations of Freedman group **(Figure 1G)** (19). In contrast to IL-4 and IL-13, the level of IL-10 was enhanced significantly in murine Th2 cells treated with calcitriol, as validated by flow cytometry, qPCR and ELISA **(Figure 1E, G, I)** We also assessed IL-9 and IFN-γ levels, cytokines predominantly secreted by Th9 and Th1 cells, respectively. We found no significant changes when Th2 cells are treated with calcitriol (**Figure 1E, F, H**). This indicates that post calcitriol treatment, any alteration of Th2-specific cytokine is not because of switching to enhanced cytokine production emanating from either Th1 or Th9 cells. Concurrently, human Th2 cells treated with calcitriol led to attenuated IL-4 and IL-13 production (**Figure 2A, B**).

**Fig. 1.**
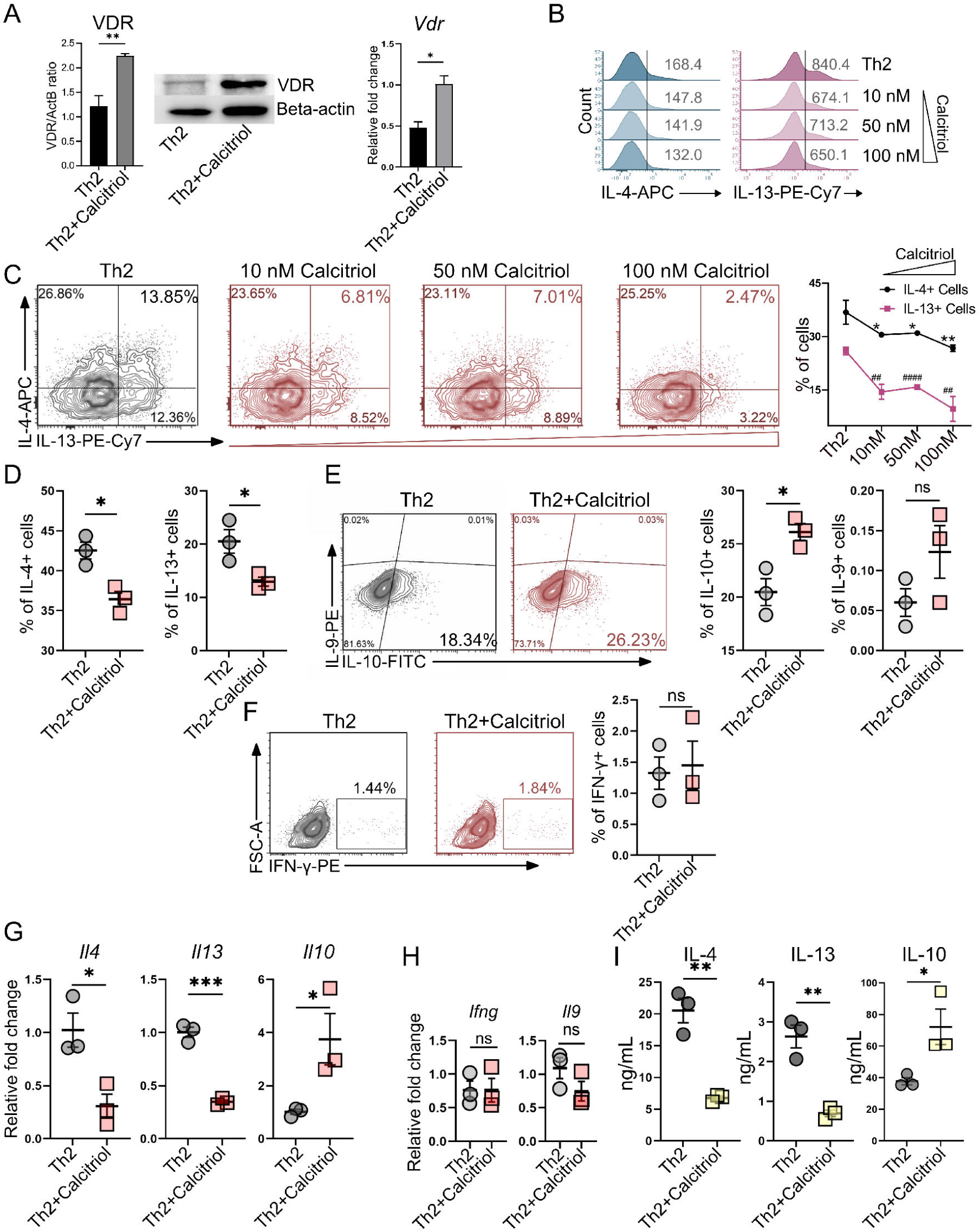
Attenuated Th2 cell differentiation post calcitriol treatment. **A**. The expression of VDR in Th2 cells treated with calcitriol has been assessed by immunoblotting (left) and qRT-PCR (right). **B, C**. The mean fluorescent intensity (MFI) and the percentage of IL-4 and IL-13-secreting cells during Th2 polarization in the presence of dose-dependent calcitriol treatment has been assessed by flow cytometry. The geometric mean of the MFI values has been shown here. **D**. The percentage of IL-4 and IL-13 secreting cells has been shown by flow cytometry post calcitriol treatment. In this and subsequent experiments, 100 nM of calcitriol has been used during differentiation. **E, F**. The percentage of IL-9 and IL-10-secreting cells (**E**) and IFN-*γ*-secreting cells has been assessed in untreated Th2 cells or Th2 cells treated with calcitriol. **G, H**. The expression levels of *Il4, Il13, Il10* (**G**) and *Ifng, Il9* (**H**) genes have been measured by qRT-PCR when Th2 cells are treated with calcitriol. **I**. The amount of secreted IL-4, IL-13 and IL-10 is measured by ELISA from the cell-free supernatant of Th2 cells treated with calcitriol. Polarization of Th2 cells with or without calcitriol takes 5 days of in vitro culture. Each closed circle or closed square in the bars represent the mean of one experiment with error bars representing standard error of mean (SEM) of minimum three independent experiments. *p*<*0.05, **p*<*0.01, ***p*<*0.001 for changes in IL-4 compared to control; ##p*<*0.01, ####p*<*0.0001 for changes in IL-13 compared to control; ns = not significant

**Fig. 2.**
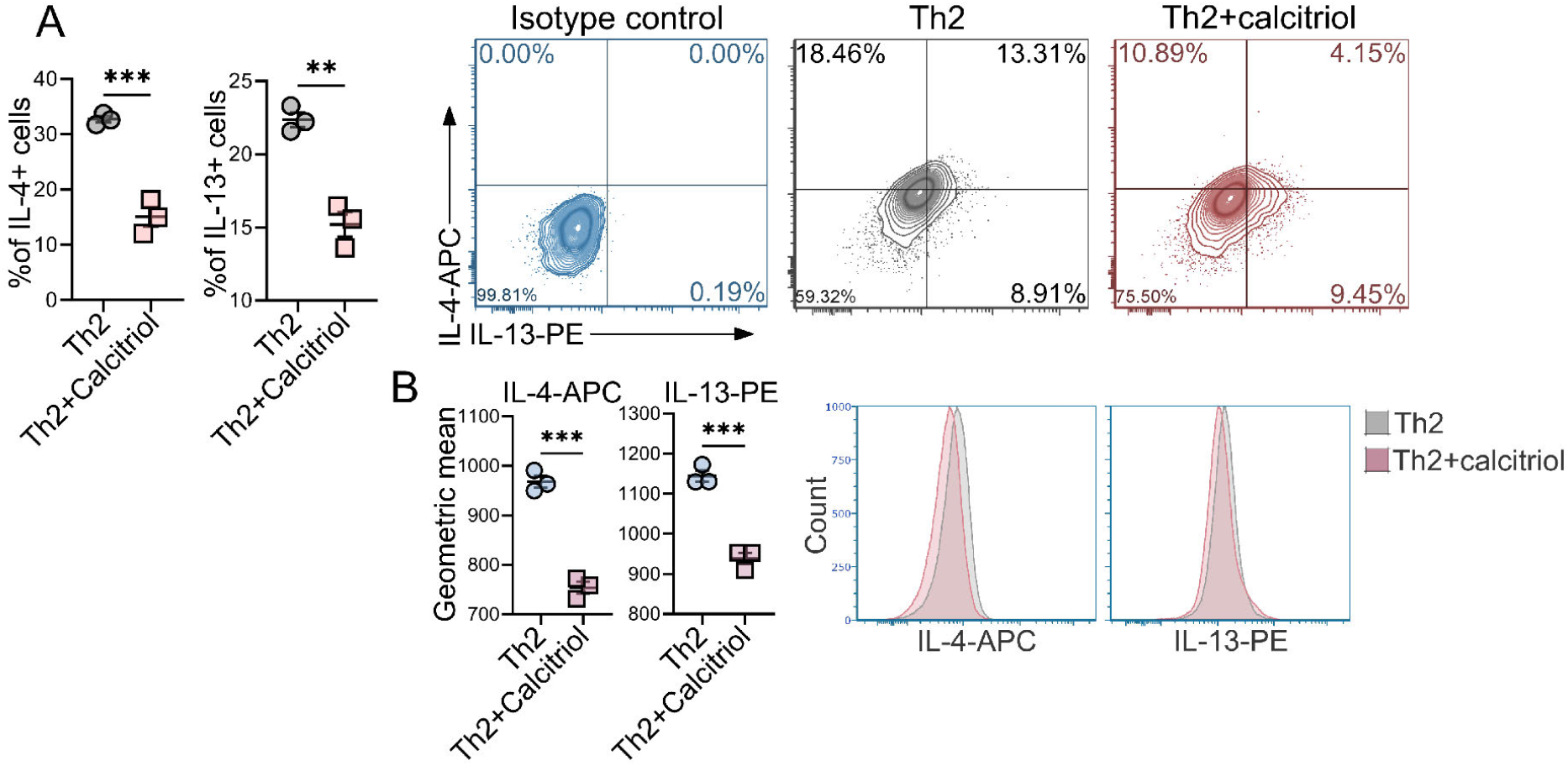
Attenuated expression of IL-4 and IL-13 in human Th2 cells post calcitriol treatment. **A**. The percentage of IL-4 and IL-13 positive human Th2 cells has been shown by flow cytometry, post calcitriol treatment (100 nM). **B**. The MFI of IL-4 and IL-13 positive human Th2 cells has also been plotted along with the bar graphs of the geometric means. *p*<*0.05, **p*<*0.01, ***p*<*0.001

### Expression of type 2 transcription factors altered post calcitriol treatment

Type 2 transcription factors (TFs) are essential for orchestrating type 2 immune response by actively maintaining Th2 phenotype (1). Gata3 is the master regulator of Th2 cells, as it suppresses Th1 phenotype. Gata3-/-CD4+ T cells failed to differentiate into Th2 cells (25). Gfi1 induced by IL-4, promotes Gata3 expression in differentiating CD4+ T cells (26). We observed calcitriol attenuating the expression of Gata3 and Gfi1 **(Figure 3A-C)**. The transcription factors Irf4, c-Maf, and Batf are also critical for Th2 cell differentiation (27). We further observed decreased expression of *Irf4* **(Figure 3A)**. Interestingly, *cMaf* level increased in calcitriol-treated Th2 cells **(Figure 3A)**. However, *Batf* expression remained unaltered in calcitriol treated Th2 cells. In our previous study, the transcription factor PU.1 expression was impaired in calcitriol-treated Th9 cells (15). We, however, observed unchanged level of *Sfpi1* (gene encoding PU.1) and PU.1 protein when Th2 cells were treated with calcitriol **(Figure 3A-C)**. Overall, our studies observed attenuation of specific but not all Th2-specific transcription factors upon calcitriol treatment.

**Fig. 3.**
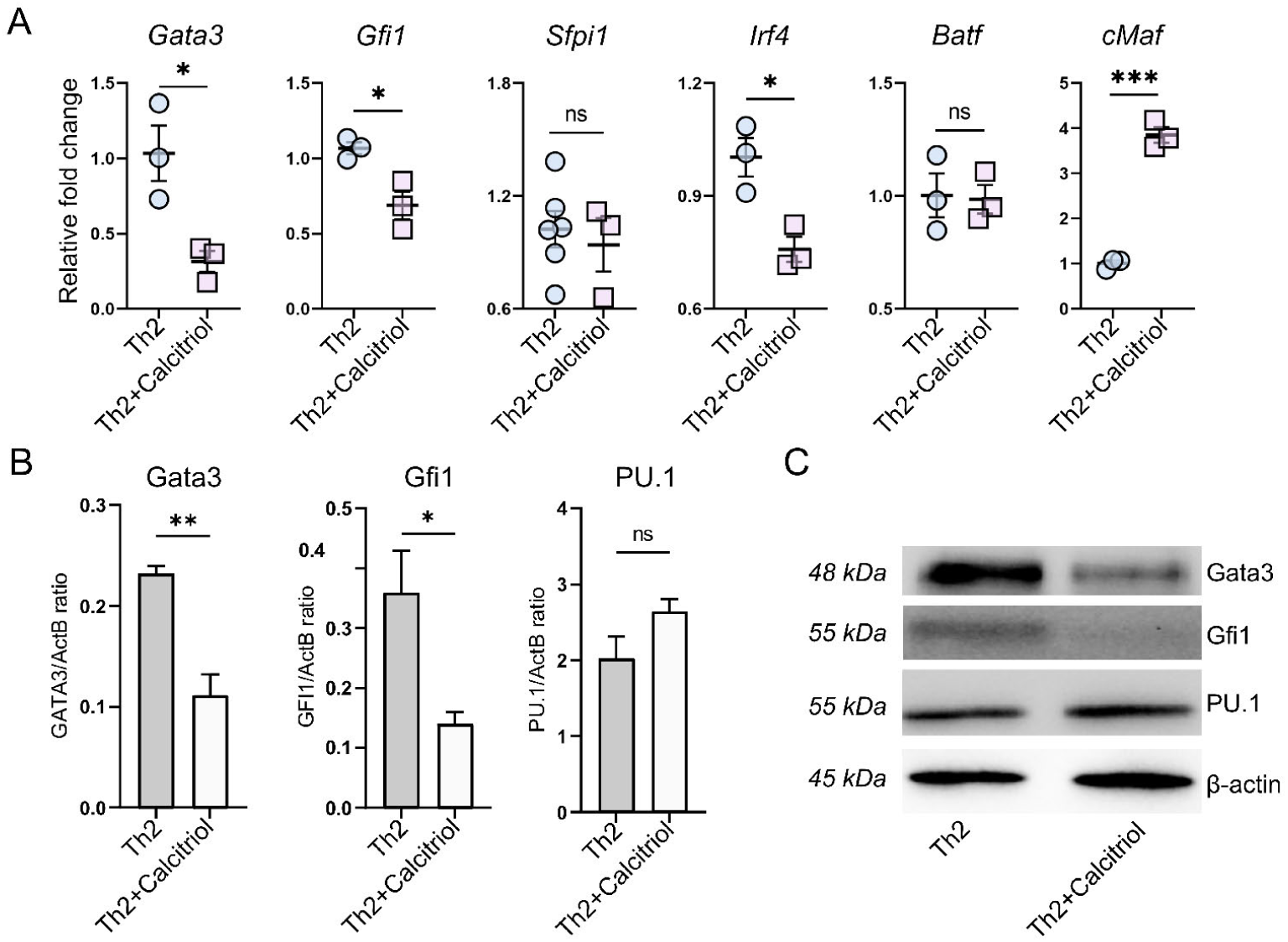
Decreased expression of Th2 cell-specific transcription factors in calcitriol-treated Th2 cells. **A**. The expression of various Th2 cell associated transcription factors Gata3, Gfi1, Sfpi1, Irf4, Batf, and cMaf are assessed in Th2 cells treated with or without calcitriol by qRT-PCR. **B, C**. The immunoblots depict the protein level of GATA3, GFI1 and PU.1 in Th2 cells treated with calcitriol compared with untreated Th2 cells. The bars represent (**B**) mean with error bars representing standard error of mean (SEM) of minimum three independent experiments. *p*<*0.05, **p*<*0.01, ***p*<*0.001, ns = not significant

**Fig. 4.**
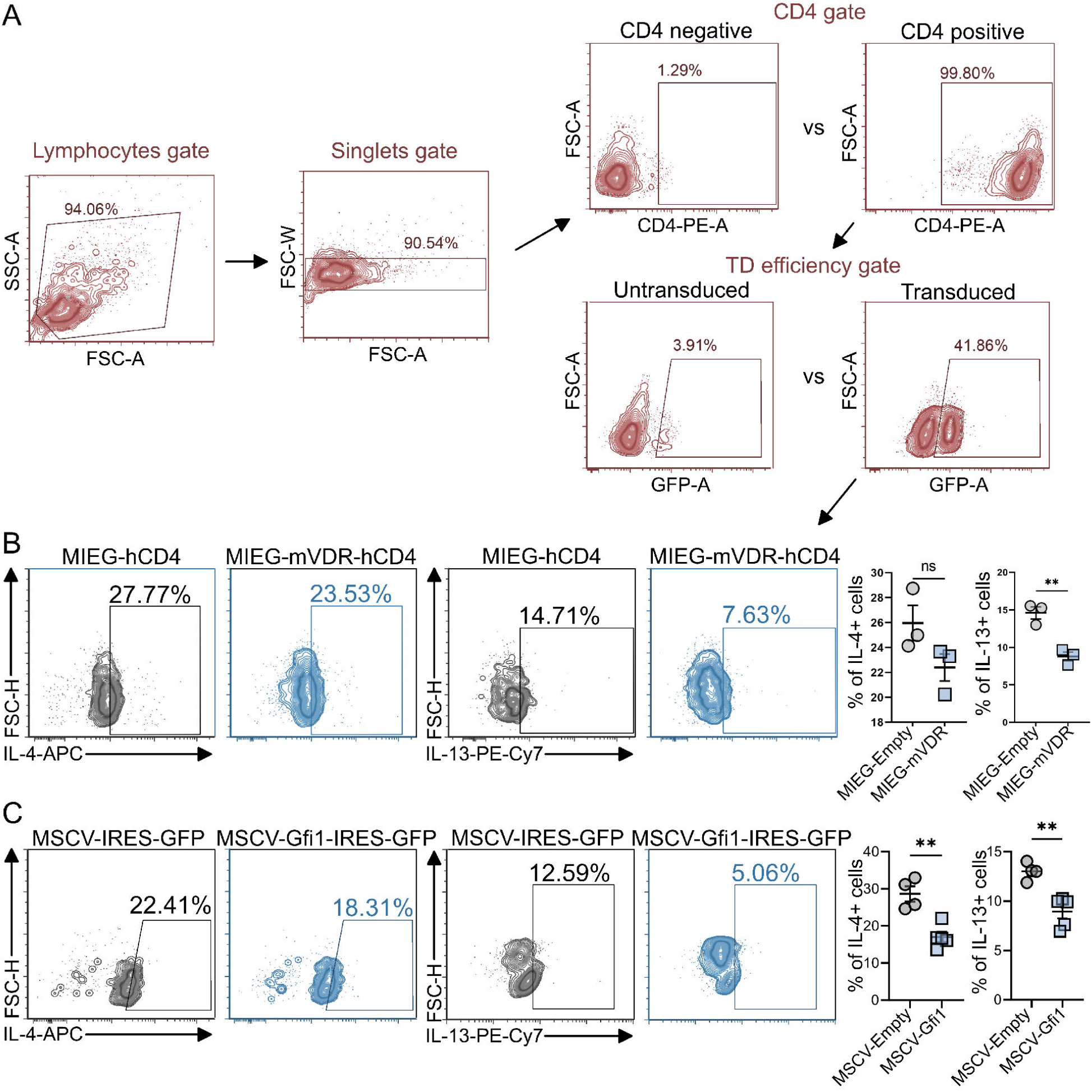
Overexpression of VDR and Gfi1 in Th2 cells have prominent effect in IL-13 production. **A**. The diagram shows the gating strategy used to identify the transduced cell population by flow cytometry. **B**. Th2 cells are transduced with empty vector (MIEG-hCD4) or with mVDR expressing vector (MIEG-mVDR-hCD4) on day 2 polarization in the presence of calcitriol. The percentage of IL-4 and IL-13-secreting cells have been assessed by flow cytometry from the transduced population. **C**. Similar to (**B**), Th2 cells are transduced with empty vector (MSCV-IRES-GFP) or with Gfi1-expressing (MSCV-mGfi1-IRES-GFP) vector in the presence of calcitriol. Shown here are the percentage of IL-4 and IL-13-secreting cells from the transduced population by flow cytometry. The bars represent mean with error bars representing standard error of mean (SEM) of minimum three independent experiments. **p<0.01, ns = not significant

### VDR directly suppressed of IL-13, but IL-4 remained unchanged; however, Gfi1 suppressed both IL-4 and IL-13 in Th2 cells

Since calcitriol increased VDR expression significantly in Th2 cells, we further wanted to elucidate the role of VDR in regulating Th2 cell differentiation. VDR was recently found to be a natural repressor for IL-13 (20). We ectopically expressed VDR in developing murine Th2 cells, and measured the percentage of IL-4- and IL-13-secreting cells. We found VDR to directly suppress IL-13 production but not IL-4, suggesting an indirect role of VDR in the regulation of IL-4 **(Figure 4B)**. The gating strategy for MIEG-mVDR-hCD4 has been shown in **supplementary figure 1A**. Previous studies have shown that Gfi1, prevented activated CD4+ T cells from triggering the T helper type 1 program (28). We used a Gfi1-expressing retroviral vector to transduce developing Th2 cells to elucidate the function of Gfi1 in Th2 cell differentiation. Th2 cells ectopically expressing Gfi1 showed a significant reduction in IL-13 and IL-4 production **(Figure 4C)**. The gating strategy to isolate transduced (for MSCV-Gfi1-IRES-GFP), CD4+ cells is depicted in **Figure 4A**. We found that ectopically expressed Gata3 substantially increased the number of both IL-4- and IL-13-producing Th2 cells (**supplementary figure 1D,E**). Calcitriol-bound VDR can interact with other TFs and form multiple regulatory complexes (14). VDR is known to interact with transcriptional repressors, such as NCOR1, COP9, and coactivators such as NCOA family members. VDR also interacts with histone acetyl transferases (HATs) and histone deacetylates (HDACs) (29). Since, the expression of Gata3 was attenuated in the presence of calcitriol, we envisaged that VDR could additionally interact with Gata3 to impair its recruitment to *Il4*/*Il13* gene locus. We observed that VDR and Gata3 co-eluted in Th2 cells in the presence of calcitriol, implying that VDR physically associates with Gata3 and thereby would potentially alter the binding of Gata3 at *Il4*/*1l13* gene **(Figure 5)**. In primary Th2 cells, we were unable to detect any interaction between Gata3 and Gfi1 (data not shown). We failed to observe VDR interaction with Gfi1 (data not shown). Whether this observation is secondary due to the lack of finding Gfi1 antibody of immunoprecipitation grade, might need to be verified.

**Fig. 5.**
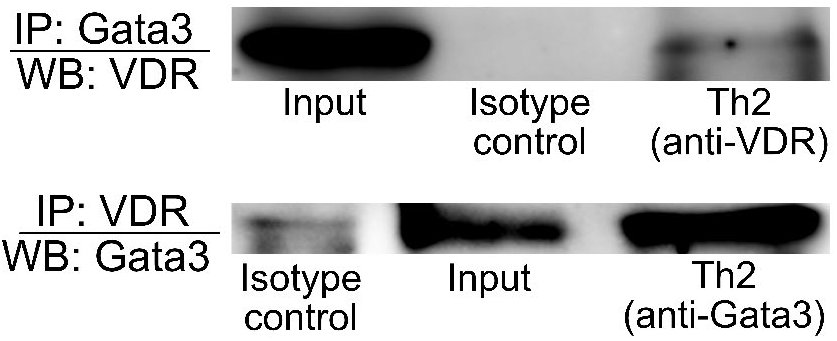
VDR interacts with Gata3 in Th2 cells. In the blot, lysates from differentiated Th2 cells are immunoprecipitated against Gata3, and the blot got developed against anti-mouse VDR (top blot). In the bottom blot in the reciprocal experiment, Th2 cells are immunoprecipitated against VDR and the blot got developed against anti-mouse Gata3.

**Fig. 6.**
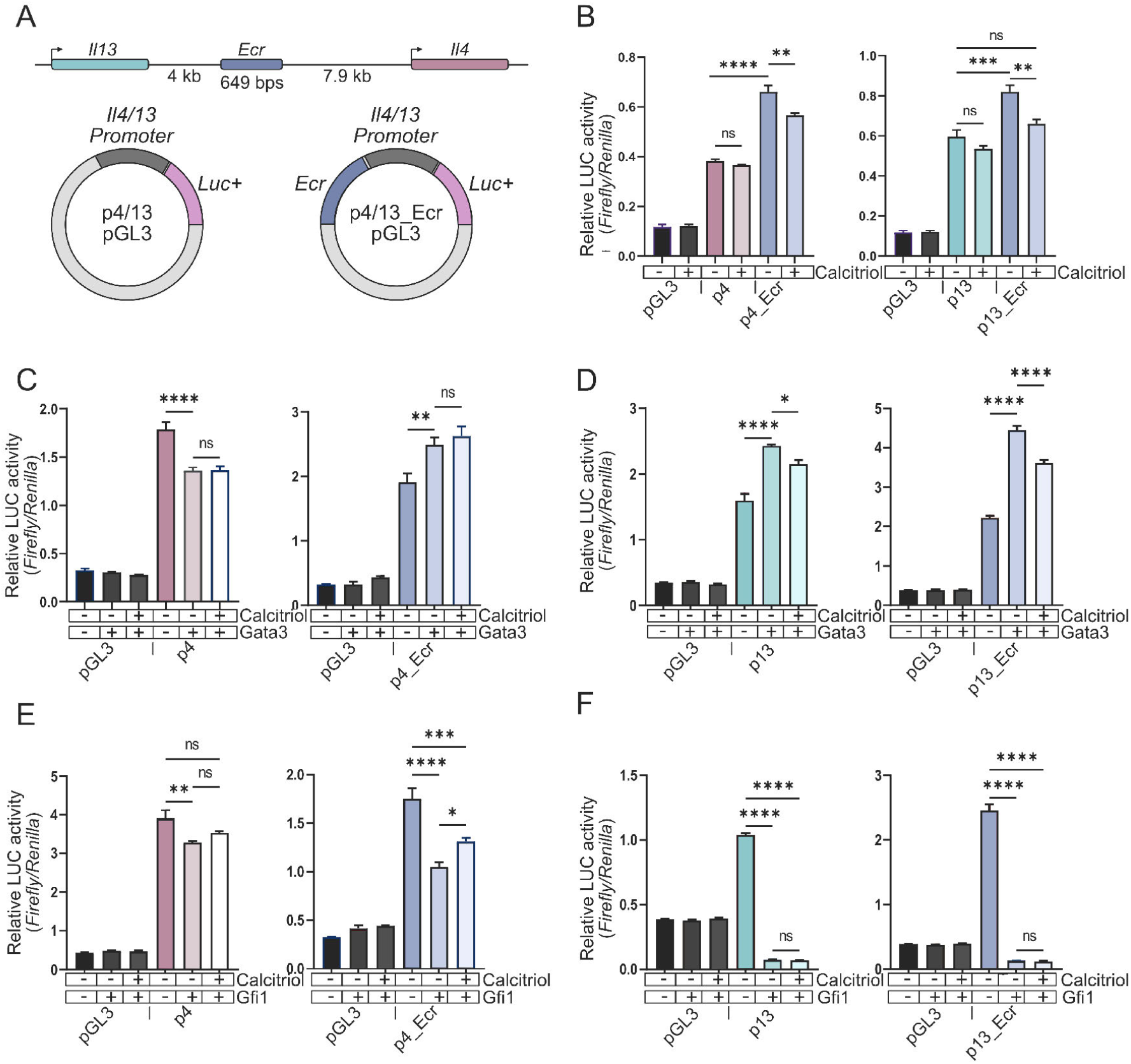
Ecr, a common regulatory element takes part in calcitriol-mediated downregulation of *Il4* and *Il13* genes. **A**. The diagram is a pictorial representation of the cloned murine *Il4* and *Il13* promoters into pGL3 reporter plasmid, with or without a common conserved regulatory element (Ecr) present between two genes. **B**. Relative luciferase activity is assessed using dual-luciferase assay when HEK293T cells are transfected with pGL3 vector only (negative control), or co-transfected with *Il4*p (p4) construct or with *Il4*p_Ecr (p4_Ecr) constructs in the presence or absence of calcitriol. **C**. HEK293T cells are co-transfected with pGL3 and *Il4*p vectors along with Gata3-expressing vector with or without calcitriol (left) or with pGL3 and *Il4*p_Ecr vectors with Gata3 in the presence or absence of calcitriol (right). Relative luciferase activity is calculated using luminometer. **D**. Similar to C, the luciferase assays are performed using either *Il13*p (p13) or *Il13*p_Ecr (p13_Ecr) vectors with or without Gata3. **E, F**. The *Il4* promoter activity (p4 and p4_Ecr) is determined when Gfi1 is co-transfected into HEK293T cells in the presence or absence of calcitriol (**E**). Similarly, the *Il13* promoter activity (p13 and p13_Ecr) is measured when Gfi1 is co-transfected in the presence or absence of calcitriol (**F**). All the cells were co-transfected with mVDR (pcDNA-mVDR). The bars represent mean with error bars representing standard error of mean (SEM) of minimum three independent experiments. **p*<*0.01, ***p*<*0.001, ****p*<*0.0001

### *Il4* and *Il13* share a common regulatory element, which is also crucial for calcitriol-mediated suppression of Th2 cells

Ecr, the regulatory element located 4 kb downstream of *Il13* gene, and 7.9 kb upstream of *Il4* gene **(Figure 6A)** was identified by aligning multiple vertebrate genomes (not shown). The 650 bp stretch of Ecr is highly conserved in vertebrate genome. We generated constructs using *Il4* and *Il13* promoters (p4, and p13 constructs lacking the Ecr, respectively), or with Ecr (p4_Ecr, and p13_Ecr respectively) to assess whether calcitriol alters the activation of *Il4* and *Il13* promoters in the absence or presence of Ecr **(Figure 6A)**. We observed that the presence of Ecr could increase the activation of either *Il4* or *Il13* promoter **(Figure 6B)**. Calcitriol could only reduce the promoter activity of both *Il4* or *Il13* only when Ecr was present **(Figure 6B)**. Nevertheless, calcitriol was able to bring back promoter activity to the baseline level in the *Il13* promoter, but not in the *Il4* promoter. **(Figure 6B)**. When Gata3 was co-expressed with *Il4* promoter, *Il4* promoter activity was found to be diminished, contrary to the general observations **(Figure 6C)** (25). However, Gata3 could transactivate *Il4* promoter only in the presence of the Ecr **(Figure 6C)**; underlying the importance of Ecr in the Gata3-mediated regulation of *Il4*. When Gata3 was co-expressed, calcitriol had no effect on controlling the activation of the *Il4* promoter, either with or without Ecr **(Figure 6C)**. In contrast, *Il13* promoter activity was significantly increased when Gata3 was present irrespective of the presence or absence of Ecr. The presence of Ecr, however, enhanced the activation even further **(Figure 6D)**. Calcitriol could suppress the *Il13* promoter activity when co-expressed with Gata3, either in the presence or absence of Ecr **(Figure 6D)**.

In HEK 293T and EL4 cells, Gfi1 significantly reduced RORγt-mediated IL-17A promoter activity in a dose-dependent manner (30). To assess whether Gfi1 imparted any regulatory role in the activation of *Il4* or *Il13* promoter, Gfi1 was co-expressed with the promoter constructs. We found Gfi1 to be a negative regulator of *Il4* promoter activity, both in the presence or absence of Ecr **(Figure 6E)**. In *Il4* promoter, calcitriol failed to impact the promoter activity when Gfi1 was transiently expressed; however, when Ecr was present, calcitriol recovered promoter activity partially **(Figure 6E)**. When Gfi1 was co-expressed along with *Il13* constructs, it resulted in a complete suppression of *Il13* promoter activity, regardless of the presence of Ecr **(Figure 6F)**. Calcitriol failed to evoke any meaningful alteration of the *Il13* promoter activity in the presence of Gfi1 **(Figure 6F)**. Overall, we found Ecr to be an enhancer element for *Il4* and *Il13*, and is important for calcitriol-mediated down regulation of these genes. Gata3 works as a positive regulator for both *Il4* and *Il13*, but needs Ecr to induce *Il4*. Calcitriol down-regulates *Il13* promoter activity in the presence of Gata3, albeit with a higher degree of repression observed for *Il13* compared to *Il4*.

To decipher the domains of VDR involved in regulating *Il4* promoter activity mutations encompassing either the ligand binding domain or DNA binding domain, were constructed (ΔDBD, Δ403-422, and Δ276-422) (**Figure 7A**). Upon performing dual luciferase assay with these mutation constructs, (with or without calcitriol) we observed significant loss in the promoter suppression by the VDR mutants (**Figure 7B**). Calcitriol slightly reduced the activity of *Il4* promoter when WT VDR was transiently expressed, and moderately up-regulated the promoter activity when VDR mutants were used. However, neither of the changes reached statistical significance (**Figure 7B**). Similar to *Il4* promoter, when VDR mutants were co-expressed along with *Il13* promoter, the promoter function significantly increased (**Figure 7C**). We observed an increasing trend in *Il13* promoter activity when calcitriol was present with deletion constructs. In ΔDBD VDR, the *Il13* promoter activity was significantly increased when calcitriol was present, highlighting the importance of VDR in suppressing *Il4* and *Il13* genes.

**Fig. 7.**
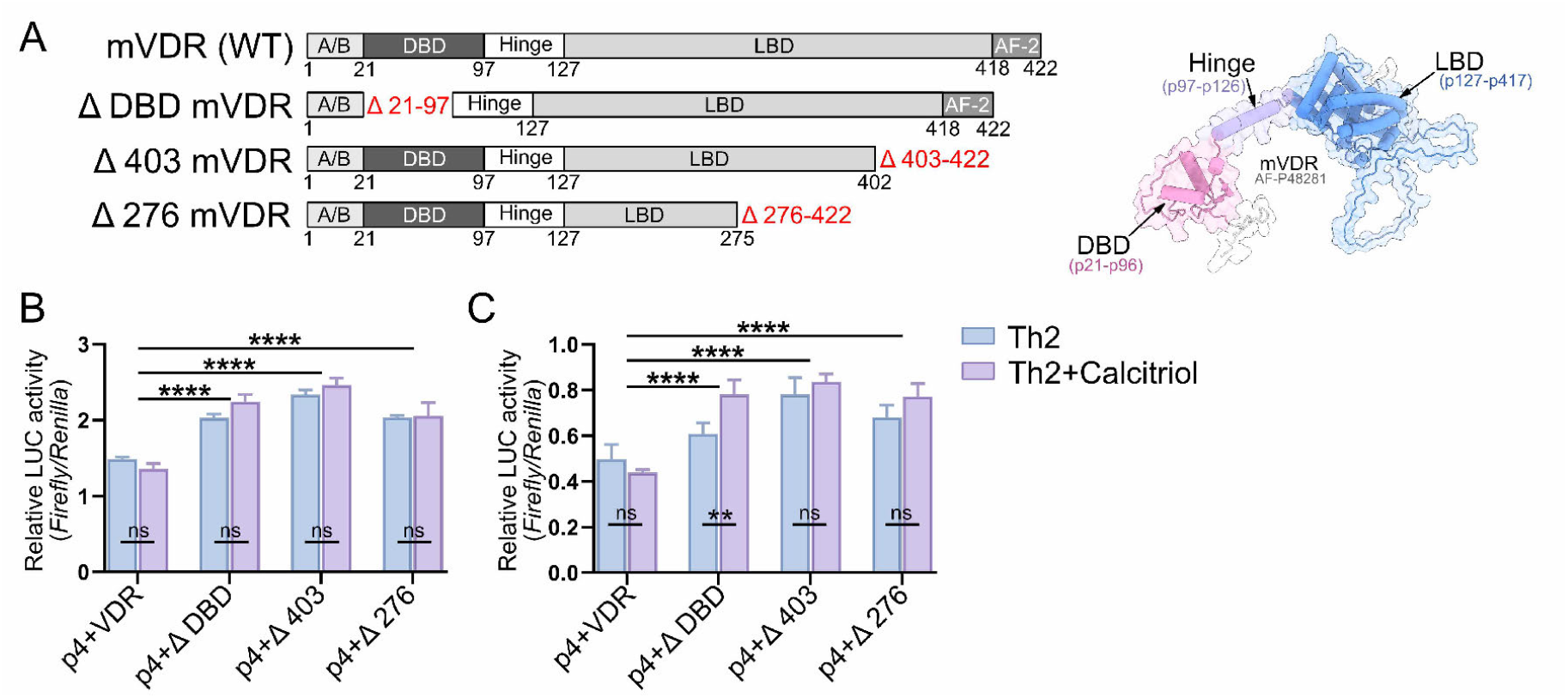
Mutations in mVDR shows prominent effect in mVDR-mediated suppression of *Il4* and *Il13* genes. **A**. The diagram depicts the structure and different domains of the mVDR protein. 21-97 amino acids (aa) forms the DNA binding domain (DBD) of the protein; whereas 127-418aa forms the ligand binding domain (LBD). Three mutation constructs of the protein have been used to establish the role of VDR in the regulation of *Il4* and *Il13* genes; a DBD deletion mutant along with two different length deletion constructs of LBD domain (Δ403-422, and Δ276-422). **B**. Relative luciferase activity is assessed using dual-luciferase assay when HEK293T cells are transfected with p4 (*Il4* promoter) in combination with the WT mVDR, and ΔDBD, Δ403 and Δ276-mVDR (pcDNA) constructs respectively, in presence or absence of calcitriol (100 nM). **C**. *Il13* promoter (p13) functions is also assessed with the same mVDR mutation constructs in presence or absence of calcitriol (100 nM). The bars represent mean, with error bars representing standard error of mean (SEM) of minimum three independent experiments. **p*<*0.01 ****p*<*0.0001, ns = not significant

### Calcitriol alters the recruitment of VDR, Gata3 and Gfi1 across different regions of *Il4* and *Il13* loci

Thus far, our research suggests that Gata3 and Gfi1 are implicated in the modulation of type 2 cytokines by calcitriol, which would result in modifications to the recruitment of transcription factors on the *Il4, Il13*, and Ecr loci. In one of our previous studies, we discovered analogous pathways utilized by calcitriol through alterations in the recruitment patterns of several TFs at the *Il9* locus in Th9 cells (15). **Figure 8A** illustrates the loci of *Il4, Il13*, and Ecr that were examined for TF recruitment via chromatin immunoprecipitation, along with the sequence and corresponding primer information. Since VDR expression was enhanced after calcitriol treatment in Th2 cells and VDR ectopic expression in Th2 cells attenuated IL-13 but not IL-4 secretion, we wanted to further determine whether calcitriol treatment altered the recruitment of VDR to *Il4*/*Il13* gene loci. By using chromatin immunoprecipitation, we were able to detect an increase in VDR recruitment following calcitriol administration, mostly at the Ecr and the *Il13* promoter, rather than at the *Il4* promoter itself **(Figure 8B)**. We also observed an increase of Gfi1 recruitment at the *Il4*, and *Il13* promoter **(Figure 8C)**, post calcitriol treatment. However, there was no change in Gfi1 recruitment at the Ecr region **(Figure 8C)**. We next elucidated whether Gata3 recruitment was altered upon calcitriol treatment in Th2 cells. A prior study has demonstrated that Gata3 may aid in the production of type 2 cytokines, which in turn triggers the development of Th2 cells (31). In the current investigation, we found that Gata3 recruitment was enhanced at *Il4*p, whereas it was dramatically decreased at *Il13*p and at Ecr regions **(Figure 8D)**. We also observed very high affinity of Gata3 towards the Ecr (Ecr2) compared to the *Il4* and *Il13* promoters and upon calcitriol treatment Gata3 recruitment was impaired at Ecr2 locus **(Figure 8D)**.

**Fig. 8.**
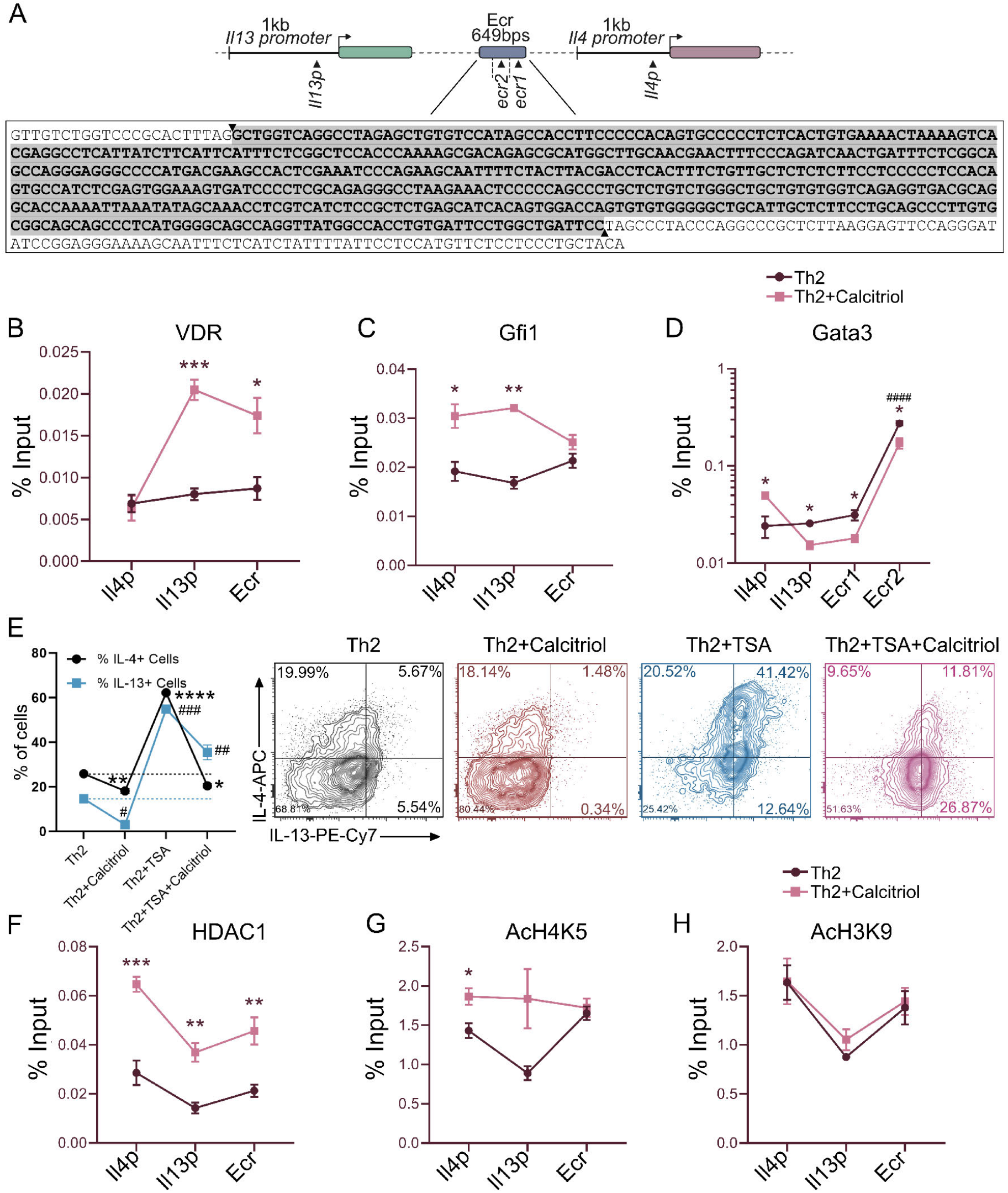
Altered histone modification and recruitment of VDR, Gata3, and Gfi1 is observed in calcitriol-treated Th2 cells. **A**. The diagram shows the relative positions of *Il4* and *Il13* promoters along with Ecr in murine genome, and its sequence, and the chromatin immunoprecipitation (ChIP) primers for the targeted positions. **B-D**. The recruitment of VDR (**B**), Gfi1 (**C**) and Gata3 (**D**) to the *Il4* promoter (*Il4*p), *Il13* promoter (*Il13*p) and Ecr region has been assessed by ChIP recruitment in Th2 cells treated with calcitriol. Signals derived from an input sample are divided by signals derived from the ChIP. The amount of chromatin used in the ChIP is represented by the input sample. qRT-PCR is used to analyze the DNA fragments. **E**. Th2 cells are treated with 100 nM calcitriol with or without 10 ng/mL Trichostatin A (TSA). After 5 days of culture, the percentage of IL-4 and IL-13-secreting cells are assessed by flow cytometry. * = changes in IL-4+ cells compared to Th2 cells; #= changes in IL-13+ cells compared to Th2 cells. One-way ANOVA is used for determining the significance. **F**. The recruitment of HDAC1 to the *Il4, Il13* promoters and the Ecr region is assessed by ChIP in Th2 cells treated with calcitriol. **G, H**. The recruitment of specific histone acetylation marks, AcH4K5 (**G**) and AcH3K9 (**H**) have been measured from ChIP experiments in Th2 cells treated with calcitriol. All the promoters and Ecr were divided into two regions, and screened separately. The data is an aggregate of those two regions. The error bars represent standard error of mean (SEM). * = difference between untreated, and calcitriol-treated Th2 samples. # = difference between immunoprecipitated regions. **p*<*0.01, ***p*<*0.001, ####p*<*0.0001

### Inhibition of HDACs resulted in an up-regulated IL-13 production, while the presence of calcitriol affected the recruitment of HDAC1 in Th2 cells

Histone deacetylase 1 (HDAC1) is a member of class I HDACs, and plays an important role in adapting to external stimuli, including stress and allergens (32). Given that Th2 cells are a major cause of allergic disorders, we anticipated that HDACs would play a part in controlling the synthesis of IL-4 and IL-13. We treated Th2 cells with Trichostatin A (TSA), which is a pan class I and II HDAC inhibitor, in the presence or absence of calcitriol. The inhibition of HDACs resulted in elevated levels of IL-4 and IL-13 expression in Th2 cells. Notably, the observed rise in IL-13+ cells was significantly greater compared to IL-4+ cells **(Figure 8E)**. Nevertheless, the administration of calcitriol resulted in a decrease in the level of IL-4 to that observed in the Th2 cells treated with calcitriol. However, it was unable to restore the levels of IL-13 **(Figure 8E)**. We also assessed the recruitment of HDAC1 in *Il4* and *l13* loci including Ecr in Th2 cells treated with calcitriol. We found an elevated HDAC1 recruitment throughout all the loci **(Figure 8F)**, indicating the importance of deacetylation in regulating type 2 cytokines. Our previous study with respect to Th9 cells, along with other studies, show how crucial histone acetylation can be to maintain distinct cytokine expression patterns in different Th cell phenotypes (15, 33). Hence, we also checked additional epigenetic markers, such as acetylated H4K5, and H3K9, along with HDAC1. We found that post calcitriol treatment, acetylation of H4K5 increased at the *Il4*p locus in Th2 cells (**Figure 8G)**. However, we could observe no changes in AcH3K9 in any locus post calcitriol treatment in Th2 cells. Our findings demonstrate the significance of epigenetic markers in regulating Th2 phenotype, as evident from the role of HDAC1 in suppressing IL-13, increased recruitment in *Il4, Il13*, and Ecr loci, and altered AcH4K5 mark in the *Il13* promoter following calcitriol treatment.

## Discussion

In this study, we have elucidated the molecular mechanisms of Th2 cell differentiation mediated by calcitriol. In contrast to the observations made by various groups, our observations are in concert with the study by Staeva-Vieira *et al*.(19). The transcriptional repressor Gfi1 was required for suppressing both *Il4* and *Il13* genes, irrespective of the presence of Ecr (**Figure 6**). The function of Gfi1 in Th2 cell differentiation has been elucidated in various studies. Th2 cell multiplication was noticeably accelerated by the forced production of Gfi1, which is brought on by T cell activation and IL-4/Stat6 signaling (34). The study observed reduced IL-4-secreting cells in Gfi1^fl/fl^-CD4Cre mice polarized under Th2-differentiating conditions. Gfi1 was crucial for controlling the IL-5 and IFN-γ production in Th2 cells (26). While this study found diminished IL-5 production in Gfi1^-/-^ Th2 cells, the difference in IL-4 production was not obvious. The discrepancy behind such contrasting observations could be attributed to the Th2 polarization protocol. Gfi1 suppressed *Ifng* gene locus activation in developing Th2 cells (26). Gfi1 has been shown to prevent alternative differentiation by Th2 cells (35).

Our study did not observe any altered expression of Th1-specific gene in calcitriol-treated Th2 cells. The down-regulation of *Il4* gene in Th2 cells by calcitriol was dependent on Gata3 (**Figure 3**). Apart from its attenuated expression in calcitriol-treated Th2 cells, Gata3 co-expression could reduce the inhibitory effect of calcitriol in mitigating transactivation of *Il4* promoter. The interaction between VDR and Gata3 in Th2 cells indicates towards possible impaired recruitment of Gata3 to *Il4* or *Il13* promoter. While the recruitment of Gata3 was significantly lower in untreated Th2 cells at the *Il4* promoter compared to calcitriol-treated Th2 cells, the recruitment was severely attenuated at the *Il13* promoter, Ecr1 and Ecr2, when Th2 cells were treated with calcitriol (**Figure 6**). Gfi1 impaired the activation of *Il4* promoter in HEK293T cells, which was partially altered in the presence of calcitriol when Ecr was present (**Figure 6**). Since GATA3 protein is stabilized by forced expression of Gfi1 in HEK293T cells, Gfi1 could play an indirect role in VDR-Gata3-mediated regulation of *Il4* gene in Th2 cells (26). It is important to note that the expression of PU.1 remained unchanged in calcitriol-treated Th2 cells, even though PU.1 interacts with Gata3 in Th2 cells (36). Therefore, distinct regulation of *Il4* promoter and *Il13* promoter was mediated by calcitriol in our studies. While IL-13 primarily causes airway hyperresponsiveness, excessive mucus production, and structural alterations in the bronchi, IL-4 stimulates both Th2 cell development and the manufacturing of immunoglobulins E (IgE) (37). While the expression of IL-13 receptor has barely been detected in lymphocytes; the expression of the receptor has been observed in murine CD4+ Th17 cells, human Th17 cells and asthma (38, 39). The fully human IgG4 monoclonal antibody Dupilumab, functions as a dual receptor antagonist of both IL-4 and IL-13 by specifically blocking IL-4R. However, the monoclonal antibody also has its fair share of side effects (40). Our study for the first time elucidates the importance of VDR-Gfi1-Gata3 axis in regulating *Il13* gene expression. Gfi1 could attenuate the activation of *Il13* promoter drastically without any additional inhibition mediated by calcitriol (**Figure 6**). Intriguingly, the recruitment of Gfi1 to both *Il4* and *Il13* promoter was observed in the presence of calcitriol (**Figure 8**). However, Gata3 recruitment was reciprocal at *Il4* and *Il13* promoters, in presence of calcitriol (**Figure 8**). Overall, these studies suggest that calcitriol-mediated severely attenuated production of IL-13, apart from IL-4 might be used as a potential adjuvant therapy.

Our results also indicate potential role of VDR-IRF4 and VDR-Batf axes in regulating Th2 cell differentiation. These findings are in concert with our previous findings where calcitriol attenuates the expression of Irf4 and Batf in regulating Th9 cell differentiation (15). The strength of TCR stimulation dictates the binding of IRF4 to VDR (41). While VDR and Batf combine to regulate the development of myeloid cells, understanding VDR-Batf axis in Th2 cell differentiation remains unexplored (42). It is possible that calcitriol mediates preferential recruitment of these transcription factors to *Il4* and *Il13* genes that could alter temporally. Since the Batf-IRF4 axis has been validated in the differentiation of Th17 cell differentiation, it would be useful to understand whether VDR-Batf-IRF4 axis is crucial for the modulation of *Il4* gene, *Il13* gene or *Il5* gene in Th2 cells (43).

Histone modification plays a key role in the expression of genes. There are histone modifications that are strongly associated with poised chromatin state dependent on cell types (44, 45). Calcitriol failed to attenuate global histone deacetylase inhibitor TSA-mediated induction of IL-13-secreting Th2 cells; however, such decrease was observed in IL-4 production (**Figure 8**). The recruitment of HDAC1 was found to be significantly enhanced only at selected locations of *Il4* and *Il13* promoter in the presence of calcitriol (**Figure 8**). Conversely, the specific histone acetylation marks, AcH3K9 and AcH4K5 remained unaltered in calcitriol-treated Th2 cells, albeit an increase in the AcH4K5 mark was observed in the presence of calcitriol only at *Il4* promoter (**Figure 8**). These observations are in contrast with our previous study involving *Il9* gene in Th9 cells (15). Since the expression of *Il9* gene remained unaltered in Th2 cells treated with calcitriol, specific histone acetylation marks were not assessed. However, our studies indicate calcitriol could act differently on different genes in similar cells. Additionally, the histone modifications could alter between distinct cytokines in the same cells. Further studies will delineate the intricate chromatin dynamics mediated by calcitriol in *Il4* and *Il13* genes that would incorporate specific histone methylation and additional histone acetylation marks.

In this study, we have proposed the detailed molecular mechanisms behind the role of calcitriol mediated regulation of Th2 cell differentiation. While the expression of *Il4* and *Il13* genes were impaired in the presence of calcitriol, the expression of *Il10* gene bucked the trend and was enhanced. We observed calcitriol altered the expression of *Il4* and *Il13* genes by a) altering the expression and recruitment of Th2-specific transcription factors (Gata3, Gfi1), b) differentially transactivating the gene promoters, c) interaction of its core receptor VDR with one of the transcription factors (Gata3) and d) distinctly regulating specific histone modifications. Additional studies will elucidate how calcitriol governs the transcription factor network in Th2 cells including Irf4 and Batf. It will also be imperative to know the domains of VDR responsible for such distinct regulation of *Il4* and *Il13* genes. Unraveling these mechanisms will go a long way to propose the potential beneficial effects of calcitriol in properly designed randomized controlled trials to treat inflammatory disorders.

## Author Contributions

**BB**: Conceptualization; Validation; Formal analysis; Investigation; Writing **SC**: Formal analysis; Investigation **SH**: Formal analysis; Investigation **RG**: Conceptualization; Resources; Writing; Supervision

## ACKNOWLEDGEMENTS

The authors acknowledge the Central Research Facility, IIT Kharagpur. The authors thank the members of the Dr. Gayatri Mukherjee’s lab, especially Mr. Debangshu Mukherjee and Ms. Debarati Biswas, for helping with flow cytometry. The authors also thank Dr. Arindam Mondal for providing help with the luminometer. Dr. Jayasree Saha from RG lab is acknowledged for critical evaluation of the manuscript.

## Supplementary Materials

**Supplementary Figure 1.**
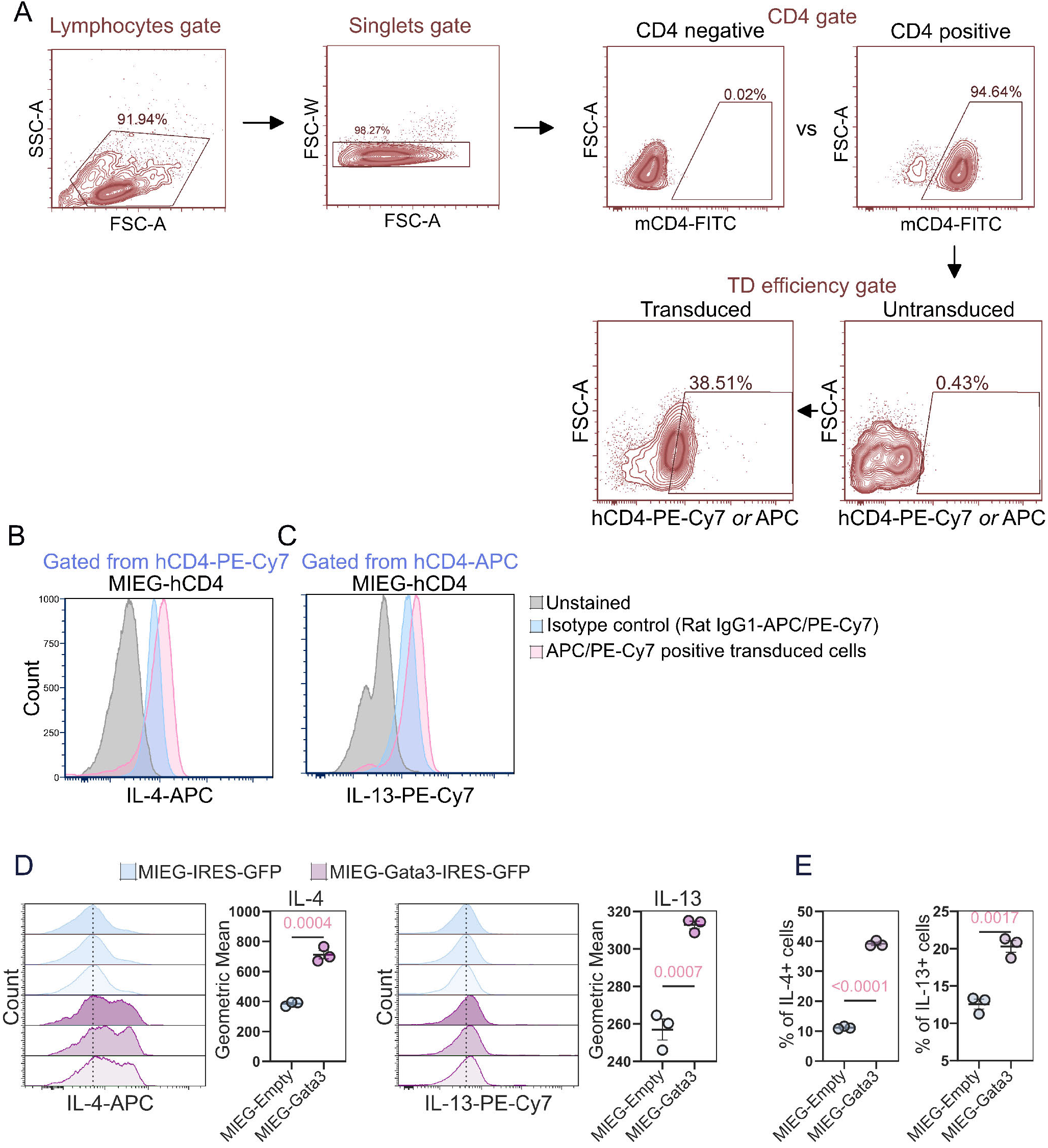
Gating strategy with isotype controls in MIEG-mVDR overexpression study, along with mGATA3 overexpression study in primary th2 cells. **A**. Gating strategy for identification o f m VDR t ransduced T h-cell p opulation v ia fl ow cy tometry. **B, C**. Th 2 ce lls tr ansduced wi th MI EG-hCD4 ex pressing retrovirus and stained with appropriate isotype controls. Rat IgG1APC is compared with IL-4 APC on the hCD4 PE-Cy7 gated cells **(B)**, and Rat IgG1PE-Cy7 is compared with IL-13 PE-Cy7 on the hCD4 APC gated cells **(C). D**. Plotted MFI as histograms of transduced Th2 cells with MIEG-IRES-GFP (empty) and MIEG-Gata3-IRES-GFP vectors. The Geometric means of three individual samples of both MIEG-IRES-GFP and MIEG-Gata3-IRES-GFP has also been plotted with a horizontal bar representing mean and error bars representing SEM, with individual values **E**. Percentage of IL-4 and IL-13 positive Th2 cells, post transduction, has also been plotted with error bars (SEM) and a horizontal bar representing the mean. The p values are mentioned at the top of each graph.

